# Spatiotemporal Complexity in the Psychotic Brain

**DOI:** 10.1101/2025.01.14.632764

**Authors:** Qiang Li, Jingyu Liu, Godfrey D. Pearlson, Jiayu Chen, Yu-Ping Wang, Jessica A. Turner, Vince D. Calhoun

## Abstract

Psychotic disorders, such as schizophrenia and bipolar disorder, pose significant diagnostic challenges with major implications on mental health. The measures of resting-state fMRI spatiotemporal complexity offer a powerful tool for identifying irregularities in brain activity. To capture global brain connectivity, we employed information-theoretic metrics, overcoming the limitations of pairwise correlation analysis approaches. This enables a more comprehensive exploration of higher-order interactions and multiscale intrinsic connectivity networks (ICNs) in the psychotic brain. In this study, we provide converging evidence suggesting that the psychotic brain exhibits states of randomness across both spatial and temporal dimensions. To further investigate these disruptions, we estimated brain network connectivity using redundancy and synergy measures, aiming to assess the integration and segregation of topological information in the psychotic brain. Our findings reveal a disruption in the balance between redundant and synergistic information, a phenomenon we term *brainquake* in this study, which highlights the instability and disorganization of brain networks in psychosis. Moreover, our exploration of higher-order topological functional connectivity reveals profound disruptions in brain information integration. Aberrant information interactions were observed across both cortical and subcortical ICNs. We specifically identified the most easily affected irregularities in the sensorimotor, visual, temporal, default mode, and fronto-parietal networks, as well as in the hippocampal and amygdalar regions, all of which showed disruptions. These findings underscore the severe impact of psychotic states on multiscale critical brain networks, suggesting a profound alteration in the brain’s complexity and organizational states.

## 1 Introduction

The human brain, with its intricate spatiotemporal organization, remains a critical focus for understanding the pathophysiology of complex mental disorders, including schizophrenia (SZ) and bipolar disorder (BP). These conditions are characterized by their diverse clinical presentations and a lack of reliable biomarkers, making accurate diagnosis and treatment particularly challenging [1–4]. Recent advancements in neuroimaging and electrophysiological research have begun to reveal that specific brain regions exhibit altered spatiotemporal dynamics that may underlie both the onset and progression of SZ and BP [5–10]. The examination of spatiotemporal dynamics, or how neural activity evolves across both time and space, offers significant promise for elucidating the neural substrates of psychosis, particularly the disruptions in brain network connectivity that are hallmark features of SZ and BP [11, 12]. Furthermore, investigating spatiotemporal complexity in brain activity has the potential to refine existing disease models, thereby enhancing diagnostic precision and informing the development of targeted, personalized therapeutic strategies [13–15]. Given the profound implications for both basic neuroscience and clinical applications, a deeper exploration of spatiotemporal brain activity in SZ and BP is essential for advancing our understanding of these complex disorders and improving patient outcomes.

When studying the human brain as a highly nonlinear, complex dynamic information processing system, it is essential to uncover the mechanisms that govern both information integration and segregation. Information integration refers to the coordination and combination of information across different brain regions, while information segregation involves the specialization of regions that process distinct types of information within its intricate networks [6, 16, 17]. The balance between these two processes is crucial for maintaining optimal brain function. Furthermore, understanding the transmission of information across various brain networks is of utmost significance, particularly given that disruptions in the integration and segregation of information are implicated in the initiation of mental illnesses [18]. Therefore, measuring and quantifying information interactions in individuals with mental disorders, as well as in healthy controls, have the potential to significantly enhance our understanding of the fundamental pathological mechanisms underlying these disorders. The proper functioning of a typical brain depends on the effective segregation and integration of regular information [19–21]. Maintaining a crucial balance between these processes is critical, as any disruption in this equilibrium could lead to severe brain disorders [22]. Identifying deviations from this equilibrium is essential for comprehending the advancement of psychotic diseases. A promising approach to gaining insights into this understanding entails quantifying information segregation and integration levels within the brain by assessing redundant and synergistic information.

Utilizing functional magnetic resonance imaging (fMRI) enables researchers to explore the intricate interplay of the brain’s functional connectivity, revealing identifiable patterns of neural activity that may provide insights into the unique characteristics of psychosis [23, 24]. Functional connectivity illuminates the intricate interplay and communication between brain regions, coordinating a variety of cognitive processes. While both low-order (pairwise) and high-order (beyond pairwise) functional connectivity are essential for understanding information exchange, recent research suggests that high-order connectivity provides a distinct advantage in revealing multi-way brain interactions [25–28]. Emerging evidence highlights the superior efficacy of high-order functional connectivity in exploring complex brain network interactions [17, 29–33]. This evolving perspective underscores the potential of advanced functional connectivity to serve as a driving force in capturing previously unknown interactions in the healthy brain and identifying novel biomarkers in the psychotic brain [34–36]. Moreover, the discovery of high-order brain networks not only deepens our understanding of neural interactions but also enhances our ability to unravel the intricate dynamics that govern the brain’s functional architecture [37, 38].

In this study, we hypothesize that the psychotic brain oscillates in an unstable, random state, with a disruption in the balance between information integration and segregation. We refer to this phenomenon “*brainquake*”. Our primary aim is to conduct a quantitative assessment of the complexities inherent in both spontaneous and psychotic brain activity. This involves with a comprehensive analysis of spatiotemporal patterns within fMRI signals. Our exploration encompasses the utilization of complexity measures, enabling us to evaluate intricate spatial and temporal patterns in brain activities. These metrics possess the capability to capture the nonlinear dynamics and high-order information encoded in spontaneous brain activity. This potentiality opens avenues for distinguishing psychotic features.

## 2 Results

### 2.1 Estimated Subject-Specific ICNs and Their Temporal Dynamics Using Psychotic rsfMRI

The data-driven brain network reference estimate was obtained through group independent component analysis (ICA) applied to a large cohort of subjects. A total of 105 intrinsic independent connectivity networks (ICNs) across multiple spatial scales (ICA model order) were identified, collectively referred to as the *Neuromark_fMRI_2*.*1_modelorder-multi template* [39], as shown in **Fig.1A** and **Fig.2**. The 105 ICNs can be categorized into 6 domains: the visual domain (VI, 12 ICNs), cerebellar domain (CB, 13 ICNs), temporal domain (TP, 13 ICNs), subcortical domain (SC, 23 ICNs), somatomotor domain (SM, 13 ICNs), and higher cognitive domain (HC, 31 ICNs), as presented in **Fig.2**.

**Figure 1:**
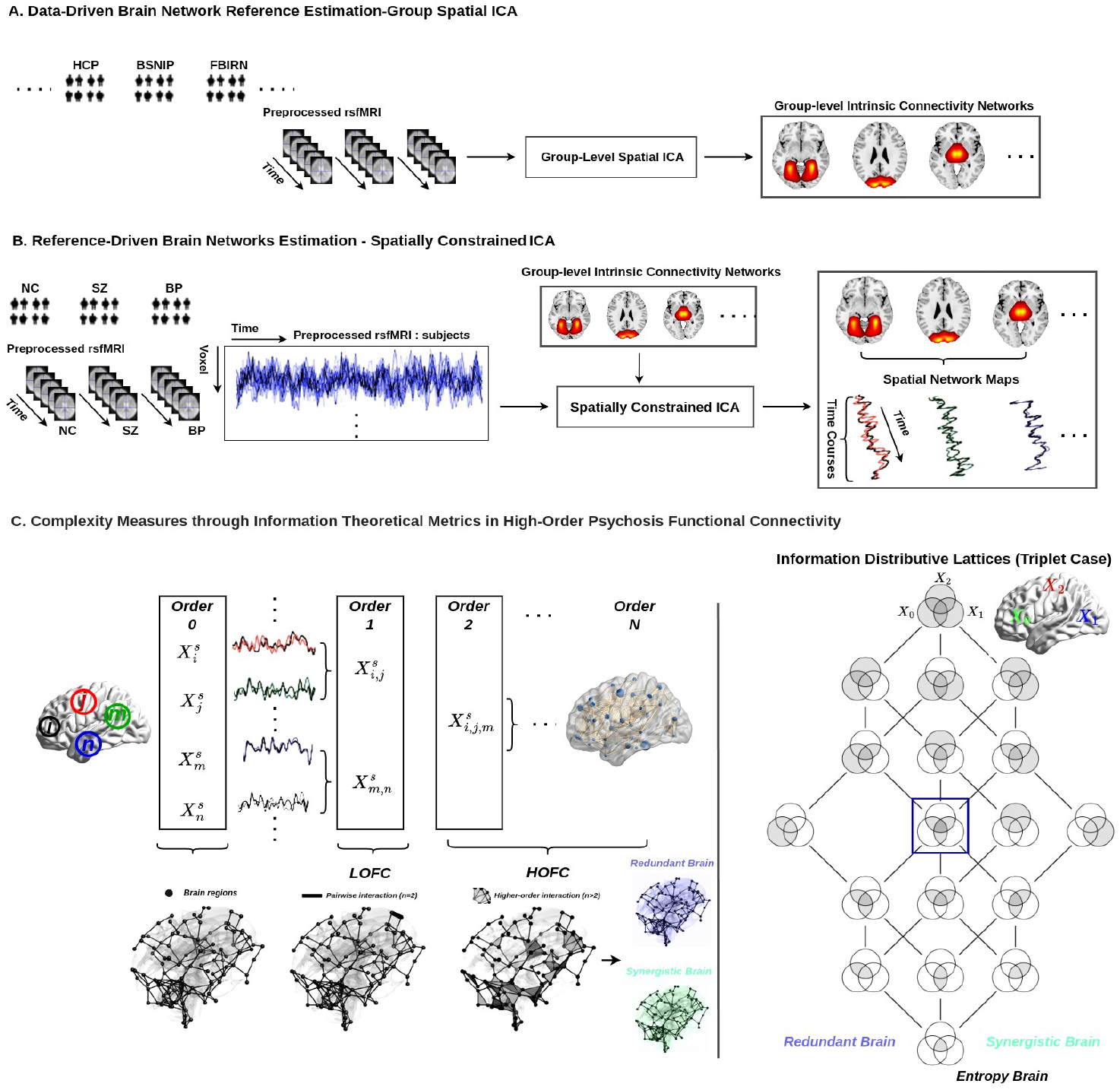
The framework designed for examining the complexity of the psychotic brain. The data-driven references for brain networks were derived from a large cohort of subjects using group-level spatial independent component analysis (ICA), as illustrated in ***A***. Spatial networks and their corresponding time courses were constructed for subjects with NC, SZ, and BP using spatially constrained ICA, based on previously established group-level intrinsic connectivity networks, as shown in ***B*** . To assess spatiotemporal complexity in the psychotic brain, complexity measures were applied to its activity, aiming to gain a better understanding of its specific spatiotemporal complexity patterns and preferences. Additionally, information interactions within brain networks are not limited to isolated (order = 0) or pairwise (order = 1) relationships. Instead, higher-order interactions (order ≥ 2) also play a significant role, highlighting the complexity of brain network dynamics, as illustrated in ***C*** . In this study, we employed integrated information decomposition, which is based on partial information decomposition (with the information distribution lattices for the triplet case presented to demonstrate the methodology for decomposing mixed information into entropy, redundant, synergistic, and total correlation components), to assess these higher-order interactions within the brain.

**Figure 2:**
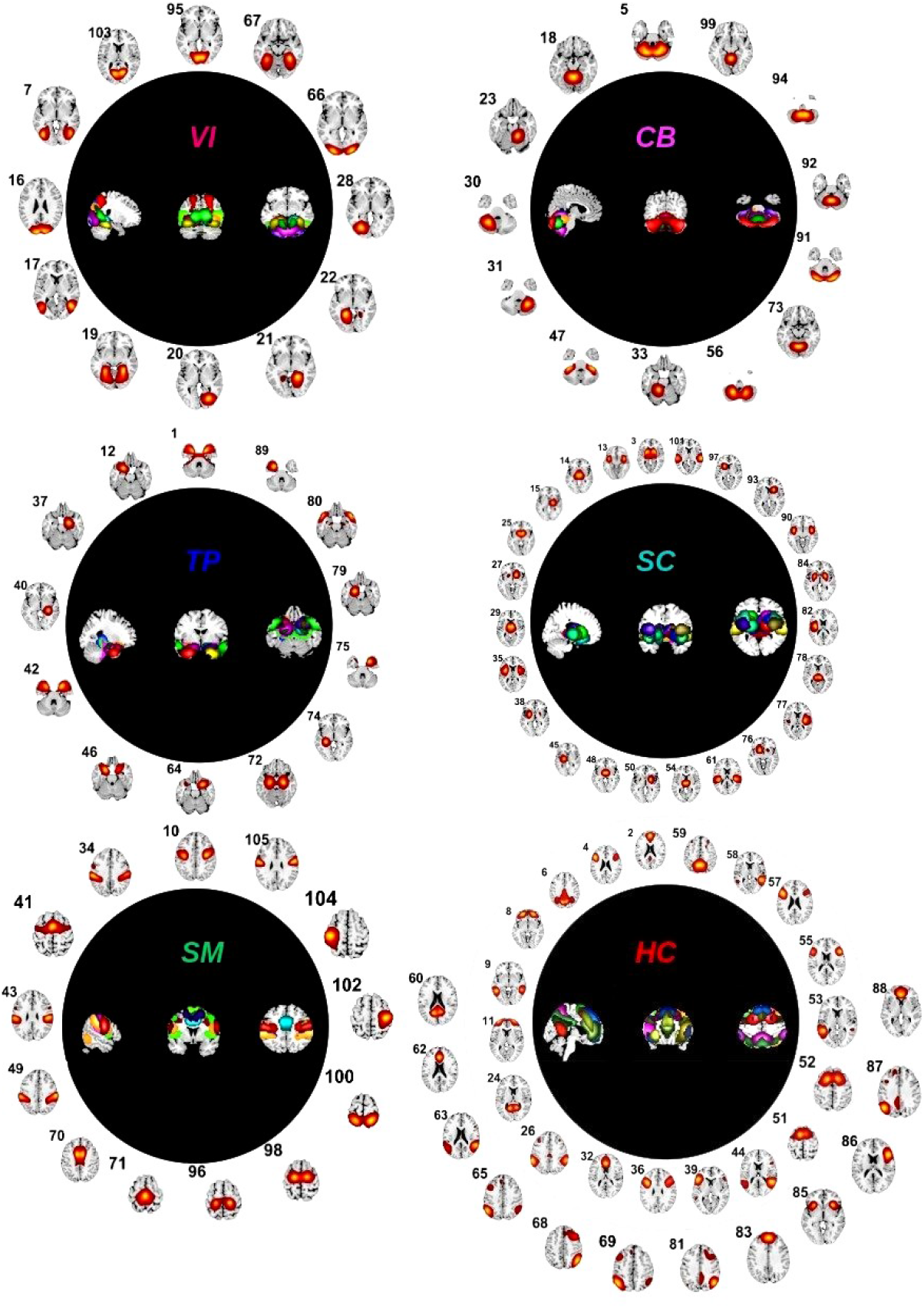
A total of 105 independent component networks were drawn from Group ICA. The *Neuromark_fMRI_2*.*1 network template* [39] is available at https://trendscenter.org/data/. It includes a total of 105 intrinsic connectivity networks (ICNs) across various domains: VI (12 ICNs), CB (13 ICNs), TP (13 ICNs), SC (23 ICNs), SM (13 ICNs), and HC (31 ICNs). The composite independent component networks are displayed in the middle section of the visual representation, providing a holistic view of their collective patterns and interactions (i.e., VI, CB, TP, SC, SM, and HC). Around this view, each individual independent component network is presented, facilitating a convenient and systematic examination of their spatial distribution and localization within the brain.

We then applied multi-objective optimization independent component analysis with reference (MOO-ICAR) to estimate subject-specific independent component networks (ICNs) and their time courses, using the *Neuromark_fMRI_2*.*1 network template* as a spatial prior brain network template. This template includes 105 high-fidelity ICNs identified from over 100K subjects [39]. The MOO-ICAR was performed on normal controls (NC), schizophrenia (SZ), and bipolar disorder (BP), and the corresponding time courses for each identified ICN were also extracted, as shown in **Fig.1B**.

After that, we applied spatiotemporal complexity measures using information-theoretical methods to quantify the state of the psychotic brain. Specifically, we used fuzzy recurrence plots and sample entropy to explore the spatial and temporal complexity of SZ and BP. We then employed integrated information decomposition, an extension of partial information decomposition (PID), to break down the information into redundancy, synergy, and unique components, enabling us to assess information integration and segregation in the psychotic brain. Following this, we applied group theory and total correlation to evaluate the high-order topological organization of information in the psychotic brain.

### 2.2 The Irregularities Present in the Spontaneous Psychotic Brain

To measure the spatial and temporal complexity of the psychotic brain, we applied fuzzy recurrence plots (FRP) [40, 41] and sample entropy [42]. To estimate the FRP, we first optimized the complexity parameter estimations, based on the minimum mutual information criterion, as shown on the left side of **Fig.3A**. The optimized parameters resulted in a delay of *τ* = 6, an embedding dimension of *e* = 5, and a tolerance of *r* = 0.081.

**Figure 3:**
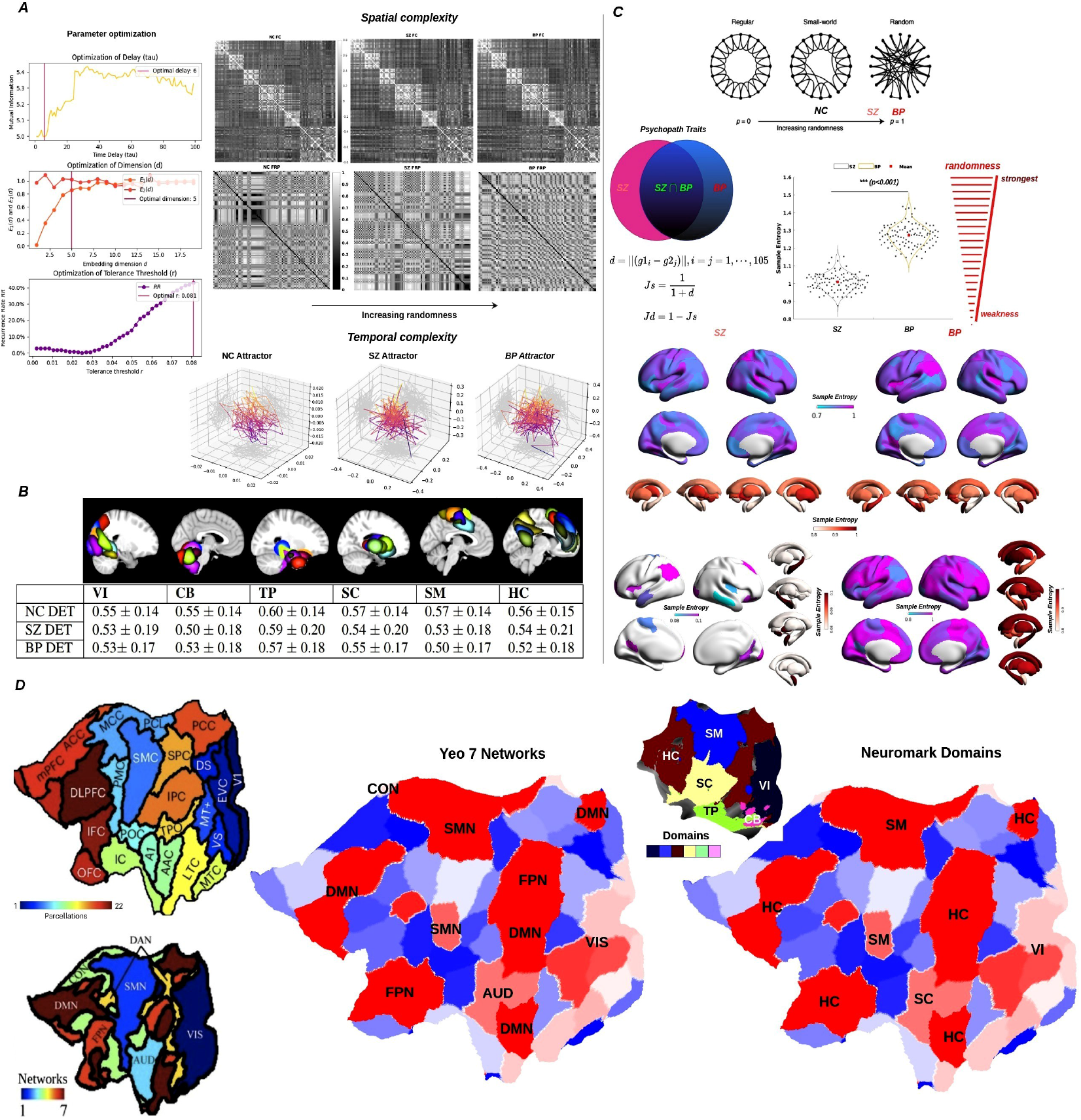
Spatiotemporal complexity measures the spontaneous brain activity in psychotic conditions. The optimized complexity parameter estimates are shown in the bottom-left corner of panel ***A***, with the following optimized values: delay *τ* = 6, embedding dimension *e* = 5, and tolerance *r* = 0.081. The functional connectivity from Pearson correlation and the FRP (optimized parameter: *e* = 3, *τ* = 1, *c* = 3) are presented in the top right side, and the related attractors for the normal controls and psychotic brain are present in the bottom in ***A***, and it shows that psychotic brain tend to randomness states. To further quantify the complexity, we applied Fuzzy Determinism (DET) to assess brain network activity, as shown in ***B*** . The psychotic brain exhibits more randomness compared to the healthy brain. According to the small-world models, studies on brain networks in SZ and BP can be categorized as favoring a less regular and more random configuration, as shown in ***C*** . The resulting sample entropy values for SZ and BP patients were projected onto the standard inflated brain surface (*fs*_*LR*.32*k*) and subcortical, revealing notable topological joint and distinctions considering SZ and BP traits are hard to separate, as shown in ***C*** . Related statistical analyses demonstrated significant sample entropy differences expressed between SZ and BP, it also suggested that psychotic brain tend to randomness state. Moreover, several brain networks associated with irregular states in SZ and BP engage multiple network systems. In panel ***D***, the color map highlights 22 functional regions [75] across the flattened left cortex (left side), with each region grouped into the Yeo 7 networks [44] as well as multiscale Neuromark_2.1 networks [39] (right side). Key networks affected in these conditions include the sensorimotor network (SMN), default mode network (DMN), central executive network (CON), frontoparietal network (FPN), visual network (VIS), and auditory network (AUD), as defined in the Yeo 7-networks. These networks correspond to specific Neuromark_2.1 domains, such as sensorimotor (SM), high-cognitive (HC), subcortical (SC), and visual (VI) domains.

Afterward, we constructed the FRP using the identified parameters: *e* = 3, *τ* = 1, and *c* = 3. As illustrated in the right side of **Fig.3A**. Our investigation revealed that the brains of individuals with psychosis exhibit greater unpredictability and irregularity compared to NC. Additionally, we constructed pairwise functional connectivity for comparison with the FRP. The use of FRP revealed more sensitive spatial patterns compared to binary functional connectivity. This finding reaffirmed our earlier conclusion that FRP stands out as an superior descriptor for functional connectivity in contrast to traditional Pearson correlation metrics [41]. Furthermore, our analysis extended to the temporal dimension, where we carefully tracked the trajectories of brain activity in individuals experiencing psychosis. The results unveiled that the psychotic brain tends to manifest increased randomness and irregularity in its activity compared to the normal brain, as illustrated in **Fig.3A**. To further quantify brain network complexity, we applied fuzzy determinism (DET) [40], a metric that assesses whether brain activity tends toward more chaotic or random behavior [41]. A small value of DET indicates that brain activity is more likely to exhibit random states. We observed that both SZ and BP exhibit randomness compared to NC, as shown in **Fig.3B**.

To more accurately quantify the state of information interaction in the brain, the small-world network is usually used as a computational model of the human brain, and this modeling approach involves initially connecting nodes with their nearest neighbors, resulting in a network characterized by a high clustering coefficient and a long characteristic path length [43]. In accordance with small-world models, investigations into brain networks in SZ and BP tend to favor a less regular and more random configuration. Here, sample entropy [42] was measured in the psychotic brain, revealing significant differences between SZ and BP. A low sample entropy suggests that brain activity in psychosis is more deterministic, while a higher value indicates increased randomness, further suggesting a tendency toward a random state in the psychotic brain. This approach is especially pertinent given the inherent difficulty in separating traits associated with SZ and BP, as illustrated in **Fig.3C**. Furthermore, we used Euclidean distance as a metric to measure their dissimilarities to improve the differentiation of important spatial topology brain regions between SZ and BP. The corresponding sample entropy values for SZ and BP were mapped onto the standard inflated brain surface (*fs*_*LR*.32*k*) and subcortical regions, illustrating notable topological similarities and distinctions. Additionally, to examine which brain domains are involved in SZ and BP, we mapped the sample entropy values onto the Yeo 7 [44] and multiscale Neuromark_2.1 templates [39], as shown in **Fig.3D**. Our analysis revealed that the most prominent irregularities and unpredictabilities in brain activity were observed in the SM, VI, DMN, FPN, TP, and SC, including regions such as the amygdala and hippocampus. These findings suggest that these specific brain domains may play a crucial role in the pathophysiology of SZ and BP, highlighting their involvement in the disrupted spatiotemporal dynamics characteristic of these psychotic disorders.

### 2.3 Disruption of Redundancy and Synergy in the Psychotic Brain

Integrated information decomposition, an extension of partial information decomposition (PID), was used to break down the group information into redundant, synergistic, and unique components [45]. The diagram of PID for triplet variables is illustrated on the left side of **Fig.4A**. Therefore, the entropy brain, redundant brain and synergistic brain can be constructed from PID [46, 47]. Meanwhile, high-order functional connectivity (HOFC) was also estimated through total correlation [25] (also known as multi-information [48] or redundancy [49]), which captures information beyond pairwise brain regions and can supply us with more rich information compared to low-order functional connectivity (LOFC) [17, 30, 34, 37].

**Figure 4:**
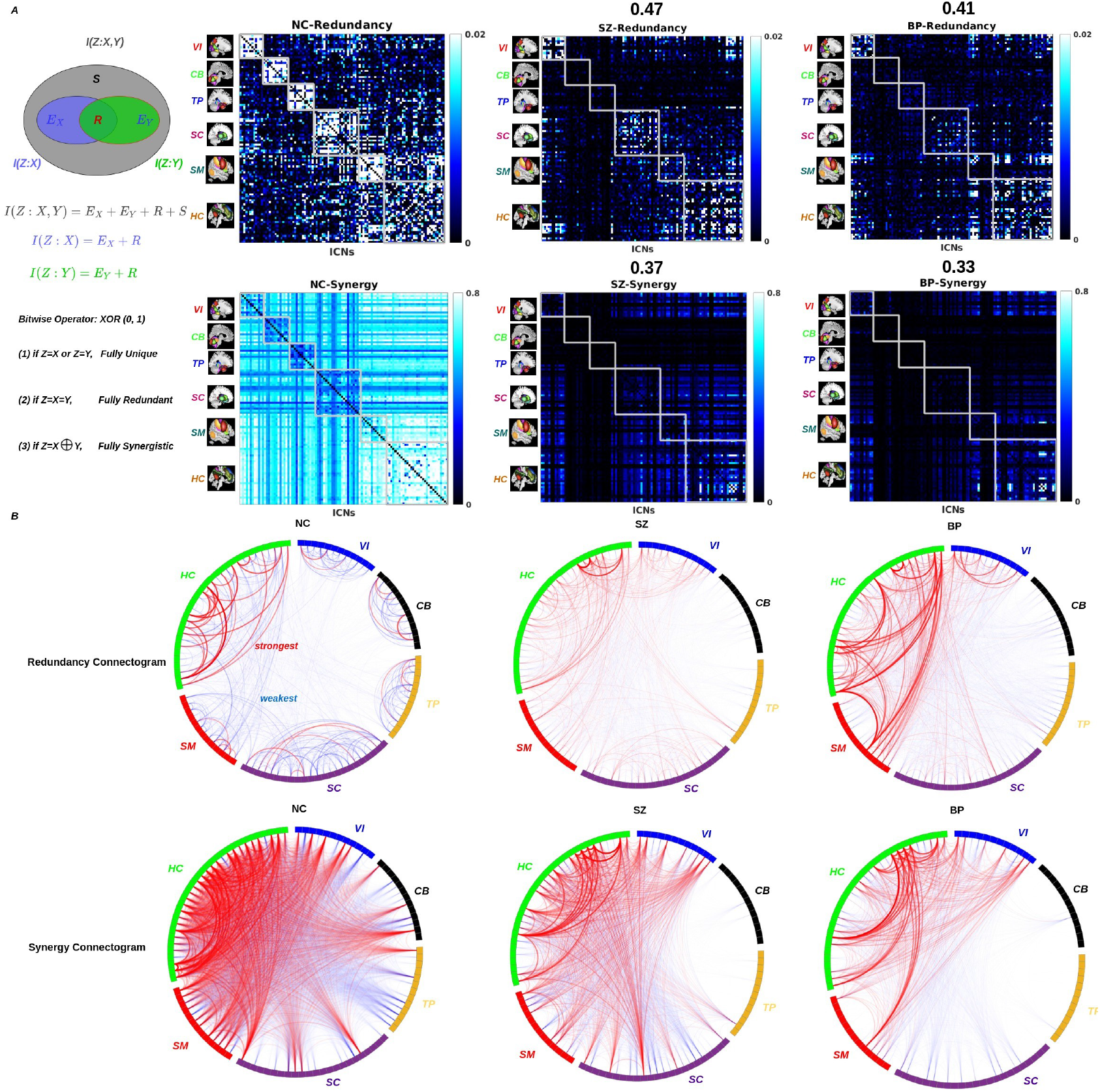
Redundant and synergistic in the spontaneous psychotic brain. The redundant and synergistic information was measured using integrated information decomposition based on partial information decomposition (PID). The basic principle of PID is illustrated in the diagram in panel ***A***. The redundant and synergistic functional connectomes between intrinsic brain domains were estimated across NC, SZ, and BP. It was observed that redundant information decreased within communities and increased between communities. Meanwhile, synergistic information gradually decreased in the psychotic brain compared to the NC. Furthermore, the correlation between psychotic redundancy/synergy and NC redundancy/synergy was labeled in the title of each psychotic matrix, respectively. The functional connectogram within the psychotic brain is shown in ***B***, depicting the redundant and synergistic connections across NC, SZ, and BP. The intensity of connections is represented by the color spectrum, with red indicating the strongest redundant and synergistic connections, and blue representing the weakest information linkages. Notably, redundant information connections primarily occurred within brain network communities in NC, whereas in SZ and BP, these connections increased between communities but decreased within communities. Additionally, a discernible pattern emerged where redundant information connections between specific brain regions were elevated, particularly between the HC and SM networks. In contrast, a reduction in redundant connections was observed between other regions, notably between the CB and TP. This intricate network analysis revealed that synergistic connections also diminished both within and between brain network communities, potentially explaining the decline in information integration observed in SZ and BP. Upon integrating both redundant and synergistic functional connectograms, our analysis suggested that the psychotic brain faces challenges in both information integration and segregation. As a result, the organization of information between brain regions appeared to adopt highly unstable or random states.

Then, using the integrated information decomposition, we constructed the redundancy and synergy of functional brain networks across NC, SZ, and BP, with the associated brain networks labeled on the y-axis, as shown in the right side of **Fig.4A**. Meanwhile, the correlation between psychotic redundancy/synergy and NC redundancy/synergy was measured as follows: *NC*_*redundancy*_ vs. *SZ*_*redundancy*_ (0.47), *NC*_*redundancy*_ vs. *BP*_*redundancy*_ (0.41), *NC*_*synergy*_ vs. *SZ*_*synergy*_ (0.37), and *NC*_*synergy*_ vs. *BP*_*synergy*_ (0.33). These results indicate that NC and SZ share a more similar functional connectivity pattern compared to BP. Simultaneously, for a more comprehensive visualization of information interaction within the brain network, the associated functional connectogram is depicted in **Fig.4B**. In this representation, the red spectrum signifies the strongest redundant or synergy connection between ICNs, while the blue spectrum indicates the weakest redundant or synergy information connection. It is evident from the results that redundant information decreases within communities and increases between communities. Simultaneously, synergistic information gradually decreases in the psychotic brain compared to the normal brain. Moreover, the major difference in redundant and synergistic functional connectivity in the psychotic brain is very clear compared to NC.

To represent the distribution of both redundancy and synergy in the human brain, we used the gradient of the synergy minus redundancy ranks. This gradient was projected onto the standard cortical and subcortical surfaces of the brain. Simultaneously, we examined the distribution of the synergy minus redundancy rank gradient across intrinsic connectivity networks (ICNs), revealing noticeable differences between normal and psychotic brains. Furthermore, the correlation between the normal brain and the psychotic brain was also measured (i.e., NC vs. SZ, r=0.19, p=0.05; NC vs. BP, r=0.20, p=0.04; and SZ vs. BP, r=0.16, p=0.09). In short summary, we showed that an unbalanced distribution between redundancy and synergy indeed happened in the psychotic brain, as shown in **Fig.5A**. Therefore, we suggested that disrupting information balance may be a major biomarker in the psychotic brain. Furthermore, we again used Euclidean distance to access their differences in order to improve the differentiation of major spatial topology brain regions between SZ and BP. The synergy minus redundancy rank gradient values for SZ and BP were projected onto the standard inflated brain surface and subcortical areas, revealing significant topological similarities and differences, as illustrated in **Fig.5B,C**. The key cortical brain domains identified in this study include the SM, VI, DMN, FPN, TP, and SC (amygdala). These findings align with the sample entropy results and suggest that the altered dynamics of redundancy and synergy contribute to the irregularity and unpredictability of brain activity, which manifest as hallmark functional brain dysregulation in SZ and BP. This abnormal balance may help explain the fragmented or disorganized thought processes observed in both disorders. By examining these disruptions in information processing, we gain valuable insights into the complex neurobiological mechanisms underlying these psychiatric conditions. Ultimately, these findings provide potential targets for therapeutic interventions aimed at restoring more typical patterns of brain network activity, which could help improve functional brain regulation in individuals with SZ and BP.

**Figure 5:**
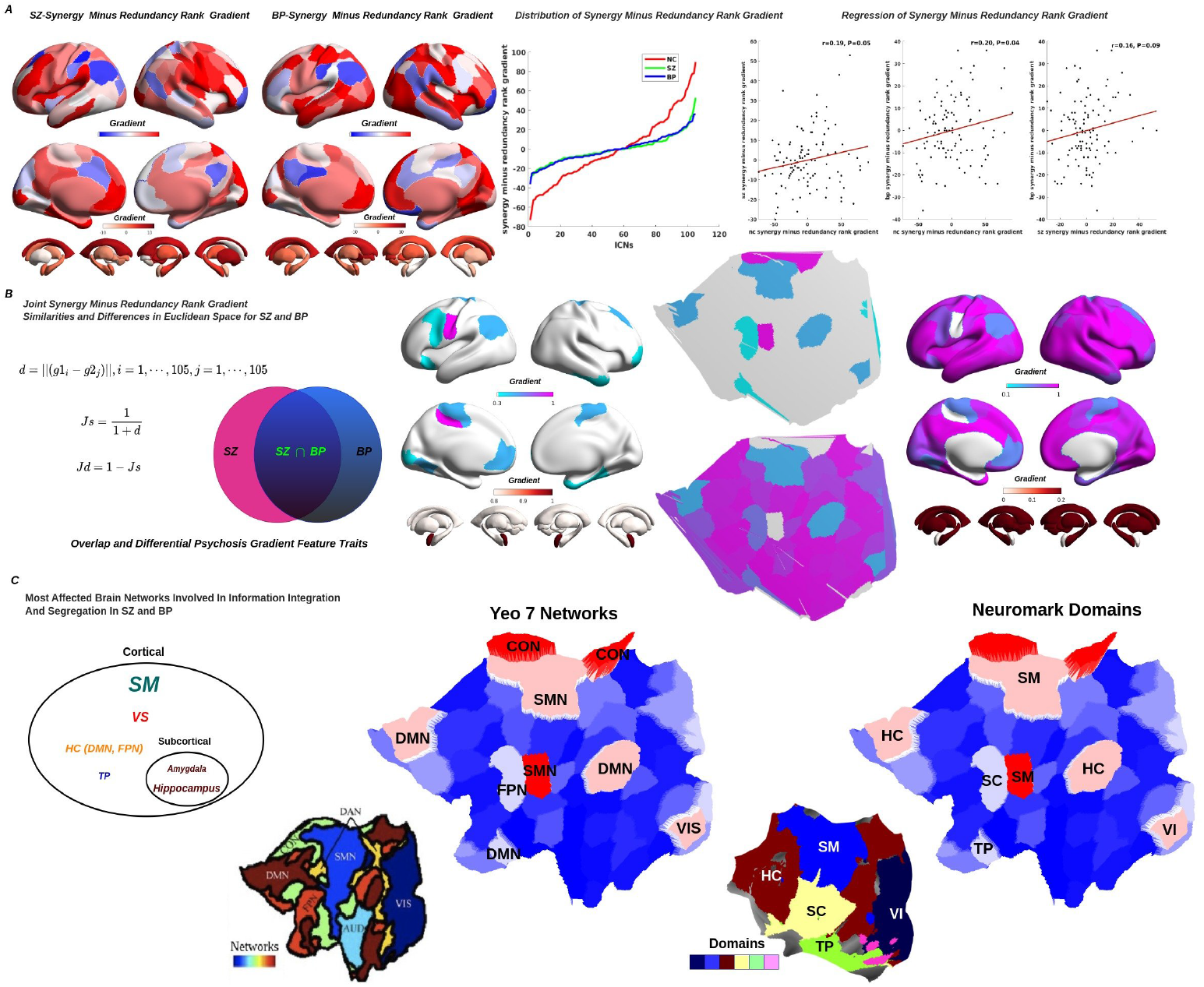
The synergy minus redundancy rank gradient in the spontaneous activity of the psychotic brain. The synergy minus redundancy rank gradient was estimated for the psychotic brain, based on redundant and synergistic interactions, and presented across both cortical and subcortical regions in SZ and BP, as shown in panel ***A***. The distribution of the synergy minus redundancy rank gradient is also presented in the middle, revealing a distinctly different gradient distribution in the psychotic brain. Furthermore, the statistical relationship between the NC synergy minus redundancy rank gradient and the SZ and BP gradients was assessed, along with the differences between SZ and BP. To further investigate the major brain domains affected in SZ and BP, we applied Euclidean distance to analyze the shared and divergent brain domains between SZ and BP across cortical and subcortical areas. Our findings revealed that the most affected brain domains included the sensorimotor (SM), visual (VS), high-cognitive (HC), and temporal (TP) domains, followed by subcortical structures such as the amygdala and hippocampus, as illustrated in panels ***B*** and ***C*** . The most affected brain networks were labeled on the flat map of the left cortex, using both the Yeo 7 and Neuromark_2.1 network templates.

### 2.4 Aberrant High-Order Dependencies in the Psychotic Brain

To explore the integration and segregation of topological information in NC and individuals with SZ and BP, we transformed redundancy and synergy functional connectivity into a tree graph layout, allowing us to visualize their topological interactions, as depicted in **Fig.6A**. Notably, we observed that SZ and BP exhibited distinct topological distribution patterns compared to NC. Additionally, individuals with SZ and BP showed greater segregation and less integration than NC subjects. This indicates reduced coordination between different brain domains in SZ and BP, reflecting less synchronized or coherent connectivity compared to NC. These disruptions in typical network organization may result in more segregated and less integrated brain activity or functional connectivity in individuals with SZ and BP.

**Figure 6:**
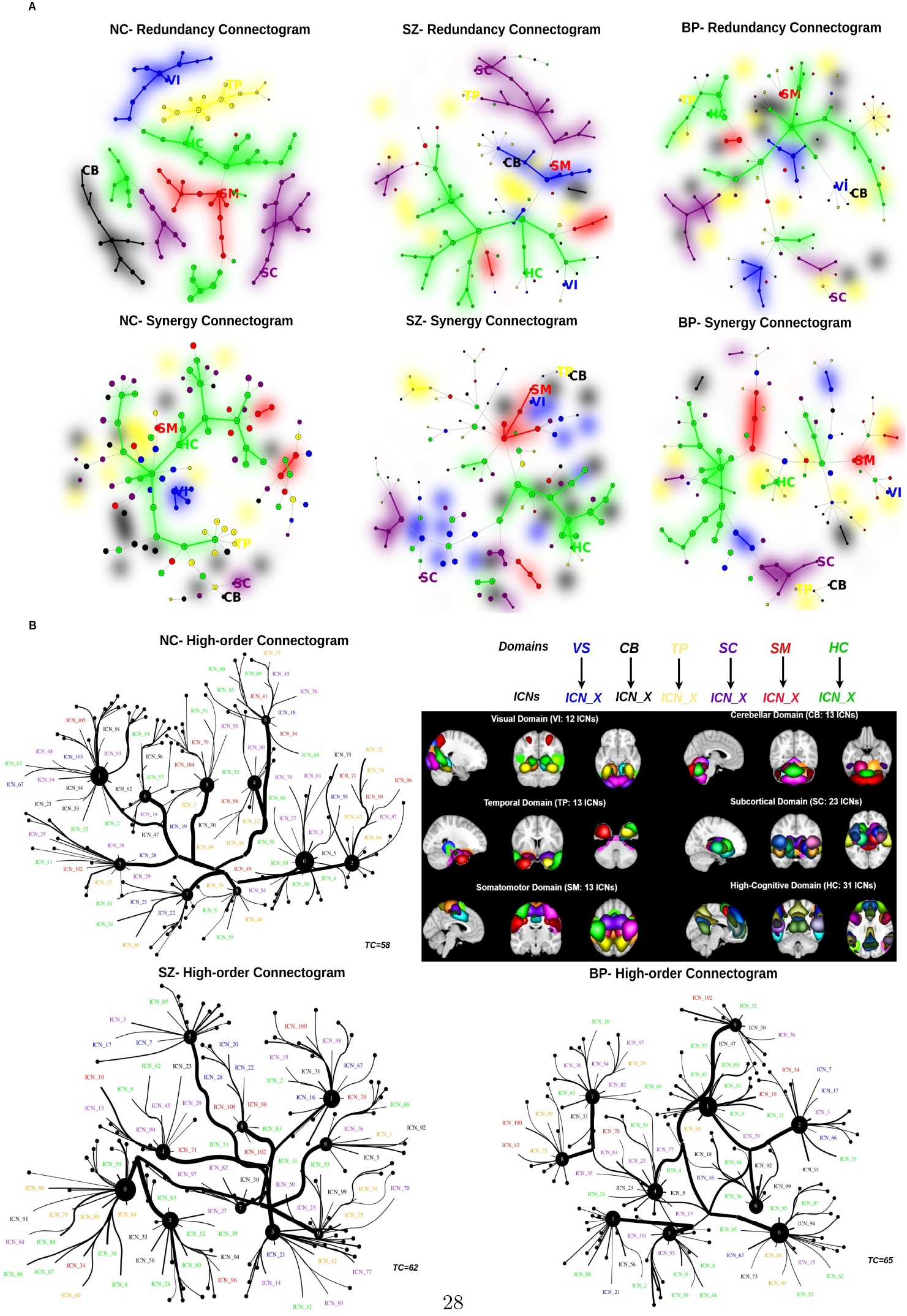
Topological functional connectome in the psychotic brain. To examine the topology of redundancy and synergy representations in NC, SZ, and BP, we plotted the connectograms of redundancy and synergy for each group as tree graphs. The intrinsic connectivity networks (ICNs) were represented as leaves on the tree, with the color spectrum corresponding to specific brain domains. Our analysis revealed that SZ and BP exhibited disrupted topological connection structures in the redundancy connectogram compared to NC, highlighting significant differences in the organization of brain networks associated with these psychotic conditions. In addition, we observed similar disruptions in the synergy connectograms, as shown in ***A***. Taken together, these findings suggest that information integration and segregation are impaired in both SZ and BP, particularly within the sensorimotor (SM), visual (VI), high-cognitive (HC), and temporal (TP) domains. These disruptions highlight specific brain domains where connectivity patterns are altered in psychotic conditions, further emphasizing the complexity of network dynamics in SZ and BP. Furthermore, we conducted an analysis of higher-order functional connectivity using information-theoretic methods based on total correlation, as shown in ***B*** with tree graphs. The edge weight represents the mutual information between ICNs and their contributions to local total correlation. The different colors of the ICNs correspond to distinct brain domains, while the solid black dot indicates the local total correlation, derived from the connected ICNs.

To capture higher-order topological information in the brain more effectively, it is necessary to apply a high-sensitivity descriptor designed for this purpose [17]. Here we applied total correlation to estimate the high-order functional connectivity. Furthermore, our previous findings have confirmed, both theoretically and empirically, that total correlation exceeds pairwise mutual information and Pearson correlation [17, 30, 34, 37]. Consequently, we employed total correlation to assess global functional connectivity, as depicted in the tree graph presented in **Fig.6B**. Each ICN is represented by a leaf of the tree and encoded with the corresponding brain network color. The solid black dots represent the local total correlation originating from connected ICNs. Notably, we observe that the HC and TP shared the strongest information, and this balance was disrupted in individuals with SZ and BP, aligning with our previously reported results [50]. Remarkably, our findings indicate that psychotic brains exhibit significantly elevated total correlation values, specifically measuring 58, 62, and 65 *nats*. This suggests a tendency toward states characterized by local randomness in the brains of individuals with psychosis compared to controls.

### 2.5 Computational Model of *Brainquake* for the Psychotic Brain

From our systematic analysis, we demonstrated that several brain networks exhibit greater instability in the psychotic brain compared to the normal brain. To further investigate this, we used a computational model to simulate the psychotic brain, as shown in **Fig.7**. The model highlights a significant increase in instability across several key networks, including the sensorimotor, visual, temporal, default mode, and fronto-parietal networks, as well as the hippocampal and amygdalar regions. Building on this, we describe this phenomenon as *Brainquake*, a concept inspired by earthquake dynamics to characterize certain unstable brain networks. Similar to active volcanoes, these networks in the psychotic brain are more prone to sudden disruptions, in contrast to the more stable networks observed in the normal brain. This analogy emphasizes the heightened vulnerability and potential for instability in the functional connectivity patterns of the psychotic brain.

**Figure 7:**
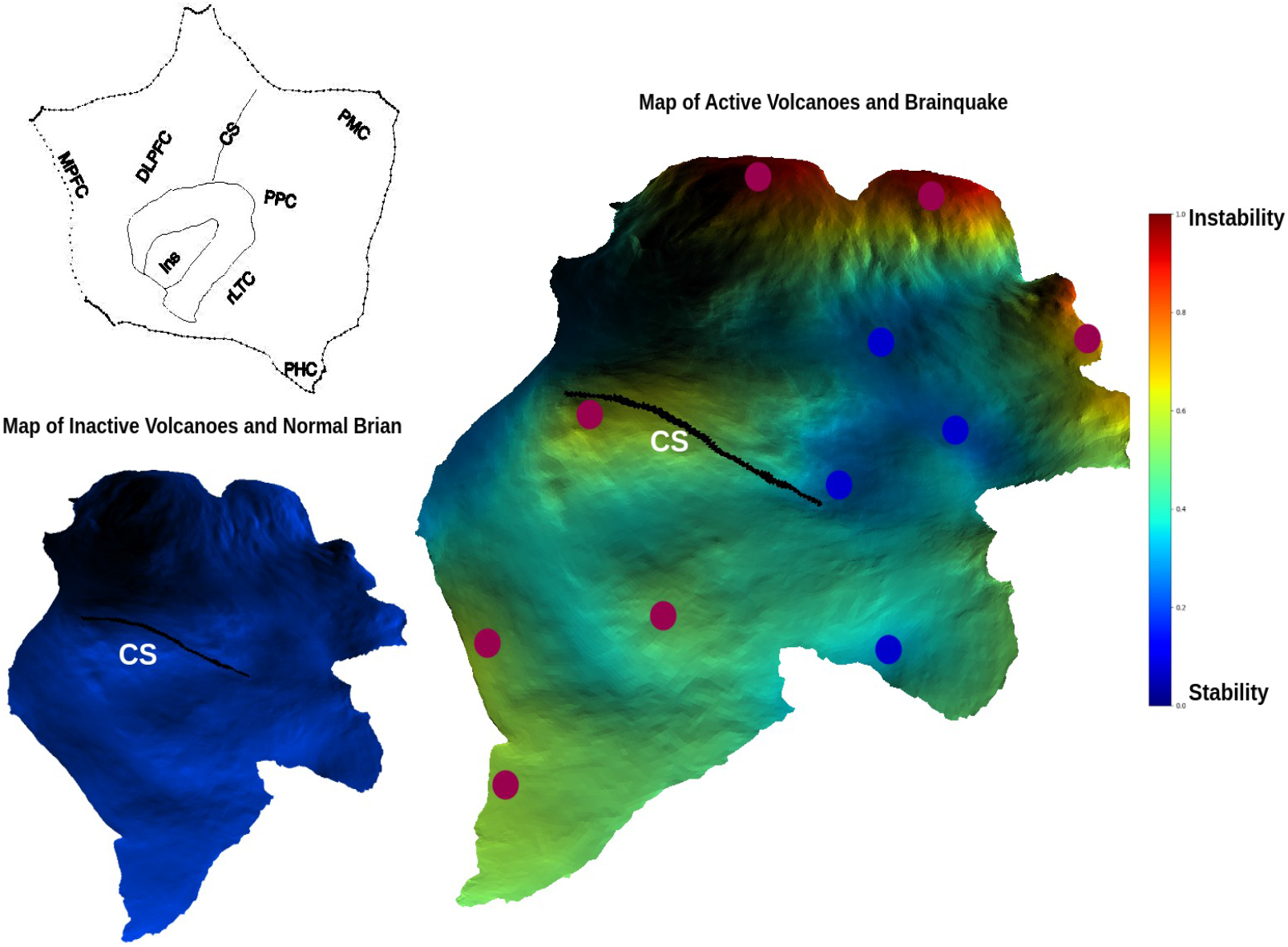
The computational model of *Brainquake* for the psychotic brain. The model of the psychotic brain suggests greater instability in several key brain networks (sensorimotor, visual, temporal, default mode, and fronto-parietal networks, as well as in the hippocampal and amygdalar regions) compared to the normal brain. The concept of *Brainquake*, inspired by earthquakes, compares certain unstable brain networks to active volcanoes, which are more prone to eruption than those in a normal brain. To aid in the recognition of brain regions on the flat brain surface, reference lines are labeled to define the inner and outer boundaries of the insula (Ins) and the central sulcus (CS). Additional key landmarks include the dorsolateral prefrontal cortex (DLPFC), posterior parietal cortex (PPC), rostral lateral temporal cortex (rLTC), posteromedial cortex (PMC), parahippocampal cortex (PHC), and medial prefrontal cortex (MPFC).

## 3 Discussion

Our spatiotemporal exploration of information-resolved brain dynamics sheds light on how the psychotic brain adapts to manage the inherent trade-off between randomness and integration. Employing group independent component analysis to estimate stable multiscale brain networks, we track spatial fuzzy recurrence plots and temporal psychosis brain activity to capture their spatiotemporal complexity. Utilizing information-theoretical metrics such as sample entropy [42] and partial information decomposition [46], we decompose the intrinsic dynamics of human BOLD signals. Through this approach, we quantify the information content carried in each brain region via sample entropy, revealing the complexity of regional dynamics. Moreover, we assess the degree to which information about the brain’s dynamics is redundantly conveyed by the current state of different brain areas, thereby illustrating their robustness. Additionally, we analyze the extent to which information is transferred across brain areas synergistically, providing insights into their integration.

We utilized fuzzy recurrence plots [40, 41] and information-theoretic metrics to assess the spatiotemporal complexity characteristics of the psychiatric brain. This assessment focused on understanding spatial and temporal neural activity patterns. Specifically, we applied complexity measures to investigate the information balance between integration and segregation within the psychotic brain, aiming to comprehend how different brain regions synchronize and desynchronize their activity. The complexity measures offer insights into the patterns of these connections, providing a means to understand how different brain regions interact and communicate under abnormal resting-state conditions [3]. In this study, we observed that the brain’s state transitions from a regular, stable state to one characterized by unstable randomness in the psychotic brain. Furthermore, we found that the delicate balance between information integration and segregation is disrupted in the psychotic brain, a phenomenon we refer to as the *brainquake*.

However, there are certain limitations to acknowledge in this study. Firstly, the assessment of the spatiotemporal complexity of the psychotic brain using fuzzy recurrence plots and sample entropy might not fully capture the predictive and regularity properties inherent in psychotic brain activity. It is essential to incorporate additional compliance measure metrics, such as the fractal dimension and correlation dimension, among others [3, 51, 52]. Integrating these metrics into the analysis would provide a more comprehensive understanding and offer additional insights into the unpredictable and irregular nature associated with diagnosing psychiatric disorders.

Secondly, for estimating redundancy and synergy from partial information decomposition, it relied on the BOLD signal matching the Gaussian distribution, and indeed, BOLD signals satisfied this hypothesis [28, 35, 50, 53]. But in order to avoid these assumptions constraints, we may need to use some universal estimators to build redundancy and synergy information [31]. Moreover, investigating bidirectional causation will uncover some causal effects in brain networks, and it may play a crucial role in understanding the psychotic brain [54, 55]. However, it is usually ignored in cause-and-effect analysis for brain networks compared to traditional cause-and-effect approaches such as Granger causality [56, 57] and transfer entropy [58].

Thirdly, dysfunction in the psychotic brain typically involves multiple functional brain networks and structural connectivity [37]. Integrating structural connectivity into our analyses can enhance the comprehension of the mechanisms underlying psychotic disorders. Furthermore, to capture a more comprehensive view of interaction information beyond pairwise brain regions, the application of high-sensitivity functional connectivity descriptors is warranted [17]. While in this study, total correlation was employed to estimate high-order functional connectivity, it is worth noting that some research suggests that dual total correlation may be superior to total correlation in capturing high-order information interactions [36]. Meanwhile, it is imperative to consider the application of simplicial complexes to model complex brain network structures because this approach enables us to capture the combinatorial properties, topology, and geometry of higher-order networks [59, 60].

Finally, to establish a connection between functional brain networks and cognitive functions, it is advisable to perform functional decoding using meta-analytic data [61]. This approach can provide valuable evidence on how psychotic brain networks are intricately linked to cognitive functions.

## 4 Materials and Methods

### 4.1 rsfMRI Dataset Acquisition

In this study, we analyzed resting-state fMRI data from 1111 subjects, including 640 normal controls (NC), 288 individuals with typical schizophrenia (SC), and 183 diagnosed with bipolar disorder (BP), all from the multi-site Bipolar and Schizophrenia Network on Intermediate Phenotypes study [62, 63]. Subjects were recruited and scanned at multiple sites: Baltimore, Boston, Chicago, Dallas, Detroit, and Hartford. The scanning period was approximately five minutes for all sites. All subjects were psychiatrically stable and on stable medication regimens at the time of the study. Participants were instructed to rest with their eyes closed and remain awake. Detailed scanning information for the entire study sample is provided elsewhere [62].

### 4.2 rsfMRI Dataset Processing

The rigorous preprocessing pipeline applied to our resting-state fMRI (rsfMRI) data, as illustrated in Fig.1, encompasses several essential steps designed to ensure the integrity and reliability of the data for subsequent analysis. First, careful quality control procedures were applied to identify and retain high-quality data, thereby ensuring the reliability of our analyses. Next, each participant’s rsfMRI data underwent a standardized preprocessing pipeline, which included rigid body motion correction, slice timing correction, and distortion correction. The preprocessed data were then registered to a common spatial template, resampled to isotropic voxels of 3mm^3^, and spatially smoothed with a Gaussian kernel having a full-width at half-maximum of 6mm.

### 4.3 Spatially constrained ICA

A spatially constrained ICA (scICA) method known as Multivariate Objective Optimization ICA with Reference (MOO-ICAR) was implemented using the GIFT software toolbox (http://trendscenter.org/software/gift). The MOO-ICAR framework estimates subject-level independent components (ICs) using existing network templates as spatial guides [5, 64, 65]. Its primary advantage lies in ensuring consistent correspondence between estimated ICs across subjects. Moreover, the scICA framework offers the flexibility to customize the network template used as a spatial reference in the ICA decomposition. This adaptability supports both disease-specific network analyses and more generalized assessments of well-established functional networks, making it suitable for diverse populations [39, 64, 66, 67].

The MOO-ICAR algorithm, which implements scICA, optimizes two objective functions: one to enhance the overall independence of the networks and another to improve the alignment of each subject-specific network with its corresponding template [64]. Both objective functions, 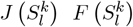, are listed in the following equation, which summarizes how the *l*^*th*^ network can be estimated for the *k*^*th*^ subject using the network template *S*_*l*_ as guidance:

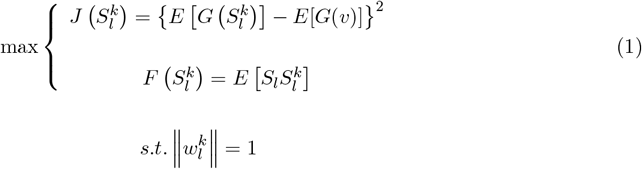

Here 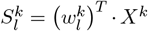 represents the estimated *l*^*th*^ network of the *k*^*th*^, *X*^*k*^ is the whitened fMRI data matrix of the *k*^*th*^ subject and 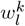 is the unmixing column vector, to be solved in the optimization functions. The function 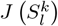 serves to optimize the independence of 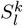 via negentropy. The *v* is a Gaussian variable with mean zero and unit variance *G*(.) is a nonquadratic function, and *E*[.] denotes the expectation of the variable. The function 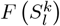 serves to optimize the correspondence between the template network *S*_*l*_ and subject network 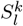.The optimization problem is addressed by combining the two objective functions through a linear weighted sum, with each weight set to 0.5. Using scICA with MOO-ICAR on each scan yields subject-specific ICNs for each of the N network templates, along with their associated time courses.

In this study, we used the *NeuroMark_fMRI_2*.*1 template* (available for download at https://trendscenter.org/data/) along with the MOO-ICAR framework for scICA on rsfMRI data. It enabled us to extract subject-specific ICNs and their associated time courses. This template includes N = 105 high-fidelity ICNs identified and reliably replicated across datasets with over 100K subjects [39]. These ICNs are categorized into 6 major functional domains: the visual domain (VI, 12 sub-networks), cerebellar domain (CB, 13 sub-networks), temporal-parietal domain (TP, 13 subnetworks), sub-cortical domain (SC, 23 sub-networks), sensorimotor domain (SM, 13 sub-networks), and high-level cognitive domain (HC, 31 sub-networks), as illustrated in Fig.2.

### 4.4 Fuzzy Recurrence Plots

Fuzzy recurrence plots (FRPs) are an advanced technique used to visualize multivariate nonlinear dynamics, particularly in the study of brain activity [41]. This approach represents data as a fuzzy cluster binary matrix, where each element reflects the recurrence of data states or phases at different time points. It can be mathematically expressed as follows:

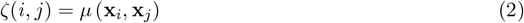

where *µ* (**x**_*i*_, **x**_*j*_) ∈ [0, 1] represents the fuzzy membership of similarity between **x**_*i*_ and **x**_*j*_. The generation of an FRP is achieved by applying the fuzzy c-means algorithm [68], which divides the dataset **X** into a collection of clusters, each denoted as *c* = 3. This algorithm assigns a fuzzy membership grade, symbolized as ζ_*ij*_ and taking values within the range of [**0, 1**] (*ζ*_*i*,*j*_ ∈ [**0, 1**]), to each data point **x**_*i*_, where *i* = 1, 2, …, *k*, concerning its association with each cluster center **v**_*j*_, where *j* = 1, 2, …, *c*. The FRP method applies the following properties:

- Reflexivity

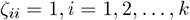
- Symmetry

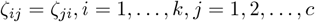
- Transitivity

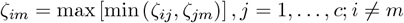

Consequently, a FRPs ∈ ℝ^*k*×*k*^ is formally characterized as a square grayscale image,

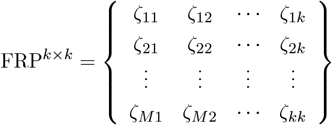

The fMRI signal can be effectively mapped into a spatial domain using a FRP, where each data point represents a state within the FRP. This transformation is especially valuable as it uncovers intrinsic patterns within complex brain activity, often providing insights that go beyond those obtainable from the original time series alone [40, 41]. Consequently, selecting the embedding dimension, denoted as *e*, is critical, as it determines the key dimensions required to capture the dynamics of brain activity. Additionally, choosing the time delay parameter, *τ*, and the number of clusters, *c*, within the time series is essential, as these parameters are aimed at optimizing the extraction of nonlinear dynamic features.

### 4.5 Dynamic Attractor

To track the temporal complexity of brain activity over time, we employed delay-coordinate embedding to construct a new vector in a reconstructed space [69]. This involved using a series of past measurements of a single scalar variable *x* from the dynamic system. The resulting *d*-dimensional vector 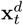 was formed from *d* time-delayed measurements of *x*_*t*_, following the procedure:

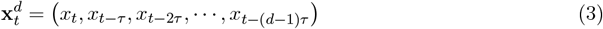

Here, we applied mutual information to optimize the embedding delay *τ* and dimension parameter *d* when utilizing the delay-coordinate embedding method. This is crucial as the values of *d* and *τ* can significantly impact the precision of trajectory estimates.

### 4.6 Sample Entropy

In dynamic systems such as the brain, brain entropy quantifies the rate at which information is generated [52]. A key complexity measure related to entropy is sample entropy (SampEn), which estimates the complexity of a system based on a known prior probabilistic distribution. SampEn has been widely used in neural time series analysis [34, 42] and is particularly useful for evaluating the complexity of brain activity in conditions such as psychosis [21]. SampEn offers two distinct advantages: it is independent of data length and is relatively straightforward to implement. For these reasons, we selected SampEn as a measure to assess the complexity of the psychotic brain.

For an fMRI signal of 𝒩 points, *U* (*i*) (1 ≤ *i* ≤ 𝒩), informally, given 𝒩 =(*u*_1_, *u*_2_, *u*_3_, · · ·, *u*_𝒩_) points with a constant time interval *t*, the family of statistics *SampEn* (*m, r*, 𝒩) is approximately equal to the negative average natural logarithm of the conditional probability that two similar sequences for *m* points remain similar. That is, within a tolerance *r* (usually refers to the distance to consider two data points as similar) and default will set to 0.2 × *std*_*U* (*i*)_, where *std* indicates standard deviation), at the next point. Thus, a low value of SampEn reflects a high degree of regularity. The parameters *m, r*, and 𝒩 must be fixed for each calculation, where *e* for embedding dimension, tolerance *r*, and number of points 𝒩 .

Let’s define a template vector with length *l*, then we have *U*_*l*_(*i*) = {*u*_*i*_, *u*_*i*+1_, *u*_*i*+2_, · · ·, *u*_*i*+*l*−1_}, and distance function, 𝒟 [*U*_*l*_(*i*), *U*_*l*_(*j*)], where *i* ≠ *j*. Then, we can define the sample entropy as follows,

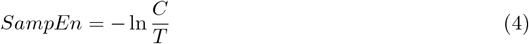

where *C* is the number of template vector pairs having 𝒟 [*U*_*l*+1_(*i*), *U*_*l*+1_(*j*)] *< r*, and *T* is the number of template vector pairs having 𝒟 [*U*_*l*_(*i*), *U*_*l*_(*j*)] *< r*. Considering properties of SampEn [42], the value of SampEn will always be zeros or positive.

### 4.7 Quantifying the Topological Similarities and Differences of the Psychotic Brain

To quantify the joint and separate topological features of brain regions between SZ and BP, we employed the Euclidean distance, Here,

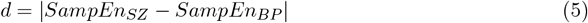

where *d* was estimated between the sample entropy, *SampEn*_*SZ*_ and *SampEn*_*BP*_ . The joint similarities were,

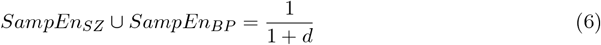

and the difference will be,

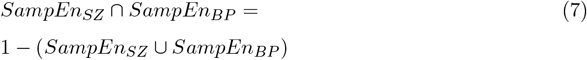

The identical approach was employed to investigate the joint and separative aspects in the topology of brain regions concerning the rank gradient of synergy minus redundancy.

### 4.8 Information Decomposition

Before discussing integrated information decomposition, we will first cover partial information decomposition (PID), as it forms the basis from which Integrated Information Decomposition is extended. The PID reveals that the two source variables *X* and *Y* given a third target variable *Z*, **I**(*X, Y* ; *Z*), can be decomposed into different types of information. That is, information provided by one source but not the other (unique information), information provided by both sources separately (redundant information), or jointly by their combination (synergistic information) [46].

In PID, the information *I*(*Z* : *X, Y*) = *E*_*X*_ + *E*_*Y*_ + *R* + *S*, where *E*_*X*_ and *E*_*Y*_ refer to unique entropy, and R and S refer to redundancy and synergy, respectively [46]. In some special cases, if *Z* = *X* or *Z* = *Y*, it explained that systems completely only have unique information; if *Z* = *X* = *Y*, it means that systems are fully redundant; and in contrast, if *Z* = *X* ⊕ *Y*, the systems are completely synergistic. Similar to PID, integrated information decomposition breaks down group information into separate components, rather than pairwise information (mutual information) as in PID, providing a direct way to decompose group information into redundancy and synergy [45]. It can be mathematically expressed as,

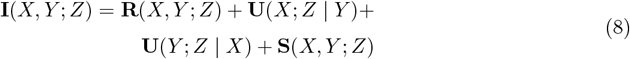

Where **R**(*X, Y* ; *Z*) refers to redundancy information, **U**(*X*; *Z* | *Y*), **U**(*Y* ; *Z* | *X*) indicates unique information, and **S**(*X, Y* ; *Z*) refers to synergy information. The PID provided us with the chance to decompose information into different pieces of information and supplied us with more information metrics to quantify the complex system, but it will present some limitations to decomposing information in the dynamic brain. Therefore, to access the redundancy and synergy in the human brain, the integrated information decomposition based on PID was introduced to estimate both metrics, considering the past states and currents of brain signal [45, 47, 70], i.e.,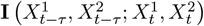, where 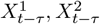 refers to past states of brain signal and 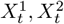 refers to current states, and redundancy, synergy, and unique information all can be avaiable from the integrated information decomposition framework. In this study, we utilized the Gaussian solver implemented in the Java Information Dynamics Toolkit (https://jlizier.github.io/jidt/) to compute all information-theoretic quantities [71].

### 4.9 Synergy minus Redundancy Rank Gradient

To assess the engagement of redundancy and synergy in the human brain, we constructed a synergy minus redundancy rank gradient, following previous studies [47]. The process involved calculating the nodal strength of each ICN across the redundancy and synergy matrix, representing the sum of all its connections in the group-averaged matrix. Subsequently, we ranked all 105 ICNs based on their connection strength, with higher-strength ICNs assigned higher ranks. By subtracting each ICN’s redundancy rank from its synergy rank, we obtained a gradient ranging from negative (indicating a higher ranking in terms of redundancy than synergy) to positive (indicating a synergy rank higher than the corresponding redundancy rank).

### 4.10 Total Correlation

Before we start to introduce total correlation, we will first introduce the Shannon entropy [72] for quantifying the information in the brain, and for the BOLD signal, consider it’s a continuous signal. Therefore, we extend the Shannon entropy to differential entropy, and for a random variable *X* with probability density function *P* (*x*) in a finite set 𝒳, the differential entropy is defined as:

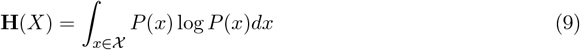

To access the information dependencies between brain regions and pairwise mutual information usually applied to capture both linear and nonlinear statistical dependencies for a pairwise BOLD signal X and Y, we have,

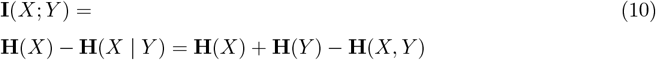

Where **H**(*X*) and **H**(*Y*) refer to the entropy of *X* and *Y*, **H**(*X* | *Y*) refers to conditional entropy, and **H**(*X, Y*) indicates the joint entropy.

Considering the limitations of mutual information [17, 30, 34, 35, 37], here we applied total correlation [25] to estimate high-order information interaction in the human brain. In general, the total correlation describes the dependence among *n* variables and can be considered a non-negative generalization of the concept of mutual information from two parties to *n* parties [25]. Suppose now we have *n* ≥ 2 variables (*X*^1^, *X*^2^, · · ·, *X*^*n*^), then the definition of TC can be denoted as follows,

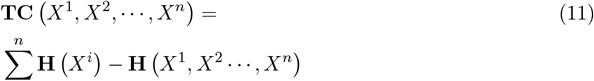

In real-life situations, estimating the marginal entropy **H** (*X*^*i*^) is straightforward, but estimating the joint entropy **H** (*X*^1^, *X*^2^ · · ·, *X*^*n*^) is considerably challenging. To address this challenge, Gaussian information theory is commonly applied to estimate total correlation because the BOLD signals satisfy Gaussian distributions [28, 35, 50, 53]. However, here we estimated the total correlation directly from the data structure itself without any prior assumptions. We use latent factor modeling to estimate total correlation, and a latent factor model is a statistical model that establishes a relationship between a set of observable variables and a set of latent variables [30, 73, 74]. The concept is to explicitly construct latent factors, denoted as *Y*, that effectively capture the dependencies present in the data. When measuring dependencies through total correlation, denoted as **TC**(*X*),

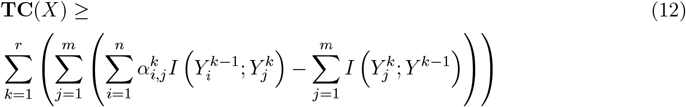

where *X* = (*X*^1^, *X*^2^, · · ·, *X*^*n*^), *Y* = (*Y* ^1^, *Y* ^2^, · · ·, *Y* ^*n*^), *m* is the number of hidden variables, *n* is the number of observed variables, *k* is the number of layers, and *α* is the learned connection matrix between the previous and current layers. Ultimately, we obtain a bound on TC that tightens as more latent factors and layers are added, allowing us to quantify the contribution of each factor to the overall bound [73, 74]. Total correlation is inherently non-negative. Through iteration optimization and a tight bound on total correlation, one can obtain the total correlation, thereby learning from the BOLD signal itself. In our study, we employed three hierarchical layers with hidden dimensions of 10, 3, and 1, respectively, to ensure that the model effectively learned all the dependencies present in the brain.

## 5 Acknowledgments

This work was supported by NSF grant 2112455, and NIH grants R01MH123610 and R01MH119251. The authors declare no conflict of interest.

## Data availability

The BSNIP data analyzed in this study cannot be shared without specific licenses. However, the dataset can be accessed through the National Institute of Mental Health Data Archive (NDA). The multi-scale order Independent Component Network (ICN) template are available online https://trendscenter.org/data/.

## Code availability

Data analysis was carried out in MATLAB version 2021a. The Java Information Dynamics Toolbox v1.5 is freely available online at https://github.com/jlizier/jidt. The ENIGMA Toolbox is available at https://github.com/MICA-MNI/ENIGMA. NeuroKit2 Toolbox is available at https://github.com/neuropsychology/NeuroKit. CorEx is available at https://github.com/gregversteeg/bio_corex. Fuzzy Recurrence Plots is available at http://www.recurrence-plot.tk/index.php. GIFT is available at http://trendscenter.org/software/gift.

## Notes

### Competing Interest Statement

The authors have declared no competing interest.

### Summary of Updates

We changed the paper's organizational style and updated the text.

